# Sun exposure drives Antarctic cryptoendolithic community structure and composition

**DOI:** 10.1101/676692

**Authors:** Claudia Coleine, Jason E. Stajich, Laura Zucconi, Silvano Onofri, Laura Selbmann

**Affiliations:** Department of Ecological and Biological Sciences (DEB), University of Tuscia, Viterbo, Italy; Department of Microbiology and Plant Pathology and Institute of Integrative Genome Biology, University of California, Riverside, CA, USA; Italian National Antarctic Museum (MNA), Mycological Section, Genoa, Italy

**Keywords:** Antarctica, Cryptoendolithic communities, Sun exposure, DGGE, qPCR, Sampling

## Abstract

The harsh environmental conditions of the ice-free regions of Continental Antarctica are considered one of the closest Martian analogues on Earth. There, rocks play a pivotal role as substratum for life and endolithism represents a primary habitat for microorganisms when external environmental conditions become incompatible with active life on rock surfaces. Due to the thermal inertia of rock, the internal airspace of lithic substratum is where microbiota find a protected and buffered microenvironment, allowing life to spread throughout these regions with extreme temperatures and low water availability. The high degree of adaptation and specialization of the endolithic communities makes them highly resistant but scarsely resilient to any external perturbation and thus, any shifts in microbial community composition may serve as early-alarm systems of environmental perturbation, including climate change.

Previous research concluded that altitude and distance from sea do not play as driving factors in shaping microbial abundance and diversity, while sun exposure was hypothesized as significant parameter influencing endolithic settlement and development. This study aims to explore our hypothesis that changes in sun exposure translate to shifts in community composition and abundances of main biological compartments (fungi, algae and bacteria) in the Antarctic cryptoendolithic communities. We performed a preliminary molecular survey, based on DGGE and qPCR tecniques, of 48 rocks with varying sun exposure, collected in Victoria Land along an altitudinal transect from 834 to 3100 m a.s.l.

Our findings demonstrate that differences in sun radiation between north and south exposure influence temperature of rocks surface, availability of water and metabolic activity and also have significant impact on community composition and microbial abundance.

## Introduction

The rate of warming due to increased levels of greenhouse gases in the atmosphere is amplified with elevation and at high latitudes due to the polar amplification phenomenon. Polar amplification predicts that as global mean temperature climbs, the greatest warming will occur at the Polar regions (Bekryaev et al., 2010). The impact of climate change is, therefore, particularly intense at the Poles and in mountain environments, nowadays known as the Third Pole (Yao et al., 2012; Yang et al., 2014). As consequence of warming, range-restricted species, particularly polar and mountain top species, have already shown severe contractions and have been the first groups in which entire species have gone extinct due to recent climate change (Parmesan, 2006; Descamps et al., 2017; Bhatta et al., 2018).

The Arctic regions are melting faster than the Antarctic and, if the heating trend continues, studies forecast an ice-free North Pole in summer by mid-century. Strong evidence of warming in Antarctica, is also documented; researchers from the British Antarctic Survey report a warming trend up to 2.5 °C since the 1940s in the Antarctic Peninsula and Maritime Antarctica, the most rapid changes in mean air temperatures on Earth (e.g. Turner et al. 2005, 2007). Previous research reported an apparent contrast between strong warming of the Antarctic Peninsula and slight cooling of the Antarctic continental interior; there are now evidences that significant warming extends well beyond the Antarctic Peninsula and covers most of West Antarctica with a warming exceeding 0.1 °C per decade over the past 50 years (Steig et al., 2009). Progressions of this warming trend will influence Antarctica’s biodiversity by the introduction of allochthonous, competitive species and the consequent extinction of highly specialized and less competitive autochthonous ones (Farrell et al., 2011; Olech and Chwedorzewska, 2011; Selbmann et al., 2012), which will have impacts on the ecosystem functions of glaciers, freshwater systems and atmosphere. Thus, it is urgent to develop a strong base knowledge for Antarctic terrestrial ecosystems and use this to identify ecosystem changes (NAS, 2011).

Endolithism is a specialized colonization by microbes to enable dwelling inside airspaces of rocks. This lifestyle represents adaptation at the edge inhabitable conditions. Airspaces within rocks offer to microbiota a protected and buffered microenvironment, allowing life to expand into different extreme conditions, i.e., hot and cold deserts or geothermal environments (Friedmann and Ocampo, 1976; Friedmann, 1982; Bell, 1993; Walker et al., 2005). Rocks are the prevailing substratum for life in the ice-free areas of Antarctica, supporting the highest standing biomass in the Antarctic ice-free desert and mountain tops emerging from the Polar Plateau (Cowan and Tow, 2004; Cary et al., 2010; Cowan et al., 2014; Selbmann et al., 2017). Endolithic microbial life represents the predominant recorded life-form in these areas (Nienow and Friedmann, 1993). The harsh conditions are considered one of the closest analogues to Mars on Earth (Quintal et al., 2018).

Different from soil microbial communities of these areas, endolithic microbes develop as very tiny and stable communities thanks to the stable and concrete nature of rocks. Various typologies have been observed for the microbial composition and the most complex and widespread are the lichen-dominated communities (Nienow and Friedmann, 1993). These self-supporting microbial ecosystems are composed of algae, mainly lichenized fungi, bacteria and cyanobacteria, many of which are endemic species to the regions (Nienow and Friedmann, 1993; Selbmann et al., 2005, 2008; Egidi et al., 2014). The high degree of adaptation and specialization in exploiting such ultimate niches makes these communities very susceptible to physical and climatic alteration (Selbmann et al., 2017) and any shift in microbial communities composition may serve as early-alarm system of environmental perturbation. Based on a substantial sampling of different typologies (volcanic and sedimentary) of colonised rocks in the Victoria Land, Antarctica, sandstone was determined to be the most suitable substratum for microbial endoliths, allowing them to spread and persist under stronger environmental pressure (Zucconi et al., 2016; Selbmann et al., 2017). To get clues to the future effects of climate change on these unique ecosystems, the response of the communities to increasing environmental pressure, due to altitude (from sea level to 3600 m a.s.l.) and sea distance (up to 100 km) was recently investigated. The results suggested that these two paramenters alone do not play as driving factors in shaping the community diversity and composition, and highlighted the needs to consider additional environmental parameters to elucidate how, in the long run, future environmental changes will impact these unique communities (Coleine et al., 2018a).

With this in mind, a new sampling campaign (Dec.2015-Jan. 2016) of sandstones was performed in the Victoria Land (Antarctica) in the frame of the Italian National Program for Antarctic Researches (PNRA), along an altitudinal gradient from 834 to 3100 m a.s.l., adding sun-exposure as new parameter to investigate its influence on endolithic settlement and development (Friedmann and Weed, 1987). Rocks with varying sun exposure (north and south faced surfaces) were collected in all the localities surveyed. Based on the prior experiences (Selbmann et al., 2017; Coleine et al., 2018a), it is suggested that sampling strategy and environmental parameters considered may have important consequences on a proper and exhaustive biodiversity description; thus, before proceeding with metabarcoding and metagenomics experiments to develop a detailed picture of microbial diversity and functionality, we have selected a suite of more rapid and cheaper complementary approaches. We have used Denaturing Gradient Gel Electrophoresis (DGGE) and quantitative PCR, to test hypothesis that changes in sun exposure impacts community composition and abundances of primary biological assemblages.

## Materials and methods

### Study area

Eight localities were visited in Victoria Land (Continental Antarctica) during the XXXI Italian Antarctic Campaign (Dec. 2015 - Jan. 2016). North and south exposed sandstone samples were collected along a latitudinal transect from 74°10’44.0’’S 162°30’53.0’’E (Mt. New Zealand, Northern Victoria Land) to 77°54’43.6’’S 161°34’39.3’’E (Knobhead, Southern Victoria Land), ranging from 834 (Battleship Promontory, Southern Victoria Land) to 3100 m a.s.l. (Mt. New Zealand) (Table 1). All rock samples were excised aseptically and collected in triplicate, transported at −20 °C at the Tuscia University (Viterbo, Italy) and stored at Mycological Section of the Italian Antarctic National Museum (MNA) until downstream analysis.

**Table 1.**
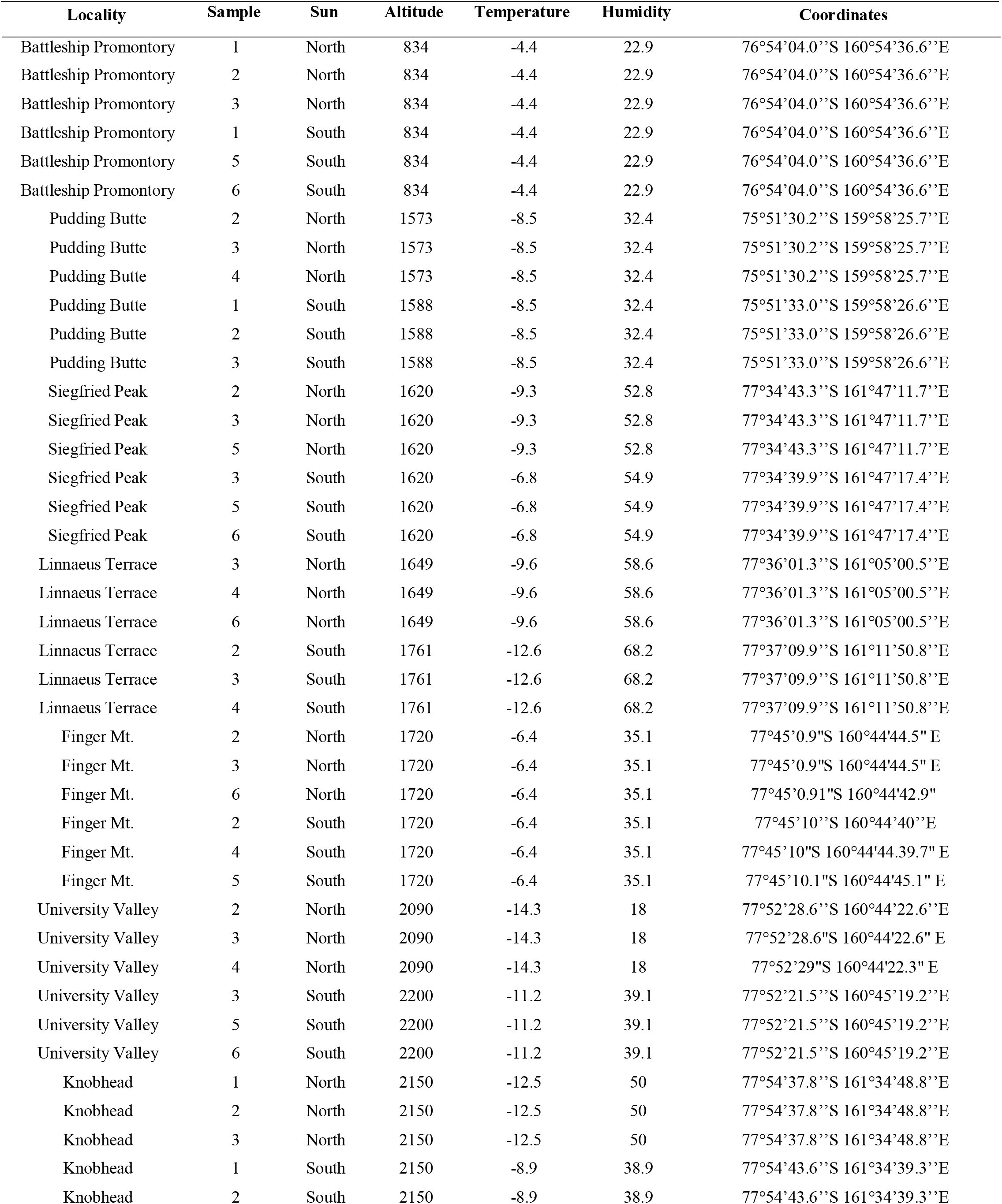
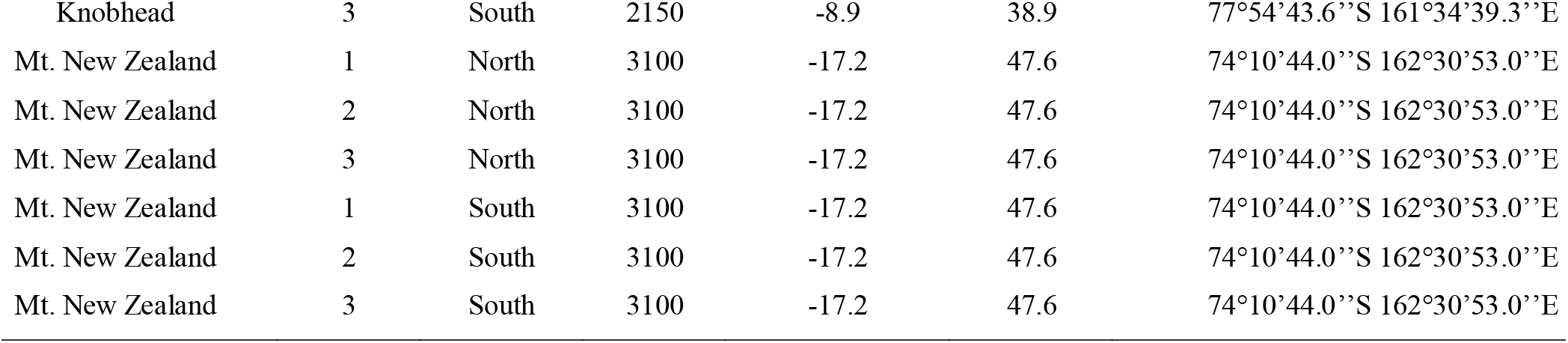
Table lists sampling details of eight visited localities in Victoria Land, Antarctica: sun exposure, altitude, air temperature (measured when sampling), relative humidity and geographic coordinates.

| Locality | Sample | Sun | Altitude | Temperature | Humidity | Coordinates |
| --- | --- | --- | --- | --- | --- | --- |
| Battleship Promontory | 1 | North | 834 | -4.4 | 22.9 | 76°54'04.0''S 160°54'36.6''E |
| Battleship Promontory | 2 | North | 834 | -4.4 | 22.9 | 76°54'04.0''S 160°54'36.6''E |
| Battleship Promontory | 3 | North | 834 | -4.4 | 22.9 | 76°54'04.0''S 160°54'36.6''E |
| Battleship Promontory | 1 | South | 834 | -4.4 | 22.9 | 76°54'04.0''S 160°54'36.6''E |
| Battleship Promontory | 5 | South | 834 | -4.4 | 22.9 | 76°54'04.0''S 160°54'36.6''E |
| Battleship Promontory | 6 | South | 834 | -4.4 | 22.9 | 76°54'04.0''S 160°54'36.6''E |
| Pudding Butte | 2 | North | 1573 | -8.5 | 32.4 | 75°51'30.2''S 159°58'25.7''E |
| Pudding Butte | 3 | North | 1573 | -8.5 | 32.4 | 75°51'30.2''S 159°58'25.7''E |
| Pudding Butte | 4 | North | 1573 | -8.5 | 32.4 | 75°51'30.2''S 159°58'25.7''E |
| Pudding Butte | 1 | South | 1588 | -8.5 | 32.4 | 75°51'33.0''S 159°58'26.6''E |
| Pudding Butte | 2 | South | 1588 | -8.5 | 32.4 | 75°51'33.0''S 159°58'26.6''E |
| Pudding Butte | 3 | South | 1588 | -8.5 | 32.4 | 75°51'33.0''S 159°58'26.6''E |
| Siegfried Peak | 2 | North | 1620 | -9.3 | 52.8 | 77°34'43.3''S 161°47'11.7''E |
| Siegfried Peak | 3 | North | 1620 | -9.3 | 52.8 | 77°34'43.3''S 161°47'11.7''E |
| Siegfried Peak | 5 | North | 1620 | -9.3 | 52.8 | 77°34'43.3''S 161°47'11.7''E |
| Siegfried Peak | 3 | South | 1620 | -6.8 | 54.9 | 77°34'39.9''S 161°47'17.4''E |
| Siegfried Peak | 5 | South | 1620 | -6.8 | 54.9 | 77°34'39.9''S 161°47'17.4''E |
| Siegfried Peak | 6 | South | 1620 | -6.8 | 54.9 | 77°34'39.9''S 161°47'17.4''E |
| Linnaeus Terrace | 3 | North | 1649 | -9.6 | 58.6 | 77°36'01.3''S 161°05'00.5''E |
| Linnaeus Terrace | 4 | North | 1649 | -9.6 | 58.6 | 77°36'01.3''S 161°05'00.5''E |
| Linnaeus Terrace | 6 | North | 1649 | -9.6 | 58.6 | 77°36'01.3''S 161°05'00.5''E |
| Linnaeus Terrace | 2 | South | 1761 | -12.6 | 68.2 | 77°37'09.9''S 161°11'50.8''E |
| Linnaeus Terrace | 3 | South | 1761 | -12.6 | 68.2 | 77°37'09.9''S 161°11'50.8''E |
| Linnaeus Terrace | 4 | South | 1761 | -12.6 | 68.2 | 77°37'09.9''S 161°11'50.8''E |
| Finger Mt. | 2 | North | 1720 | -6.4 | 35.1 | 77°45'0.9"S 160°44'44.5" E |
| Finger Mt. | 3 | North | 1720 | -6.4 | 35.1 | 77°45'0.9"S 160°44'44.5" E |
| Finger Mt. | 6 | North | 1720 | -6.4 | 35.1 | 77°45'0.91"S 160°44'42.9" |
| Finger Mt. | 2 | South | 1720 | -6.4 | 35.1 | 77°45'10''S 160°44'40''E |
| Finger Mt. | 4 | South | 1720 | -6.4 | 35.1 | 77°45'10"S 160°44'44.39.7" E |
| Finger Mt. | 5 | South | 1720 | -6.4 | 35.1 | 77°45'10.1"S 160°44'45.1" E |
| University Valley | 2 | North | 2090 | -14.3 | 18 | 77°52'28.6''S 160°44'22.6''E |
| University Valley | 3 | North | 2090 | -14.3 | 18 | 77°52'28.6"S 160°44'22.6" E |
| University Valley | 4 | North | 2090 | -14.3 | 18 | 77°52'29"S 160°44'22.3" E |
| University Valley | 3 | South | 2200 | -11.2 | 39.1 | 77°52'21.5''S 160°45'19.2''E |
| University Valley | 5 | South | 2200 | -11.2 | 39.1 | 77°52'21.5''S 160°45'19.2''E |
| University Valley | 6 | South | 2200 | -11.2 | 39.1 | 77°52'21.5''S 160°45'19.2''E |
| Knobhead | 1 | North | 2150 | -12.5 | 50 | 77°54'37.8''S 161°34'48.8''E |
| Knobhead | 2 | North | 2150 | -12.5 | 50 | 77°54'37.8''S 161°34'48.8''E |
| Knobhead | 3 | North | 2150 | -12.5 | 50 | 77°54'37.8''S 161°34'48.8''E |
| Knobhead | 1 | South | 2150 | -8.9 | 38.9 | 77°54'43.6''S 161°34'39.3''E |
| Knobhead | 2 | South | 2150 | -8.9 | 38.9 | 77°54'43.6''S 161°34'39.3''E |
| Knobhead | 3 | South | 2150 | -8.9 | 38.9 | 77°54'43.6''S 161°34'39.3''E |
| Mt. New Zealand | 1 | North | 3100 | -17.2 | 47.6 | 74°10'44.0''S 162°30'53.0''E |
| Mt. New Zealand | 2 | North | 3100 | -17.2 | 47.6 | 74°10'44.0''S 162°30'53.0''E |
| Mt. New Zealand | 3 | North | 3100 | -17.2 | 47.6 | 74°10'44.0''S 162°30'53.0''E |
| Mt. New Zealand | 1 | South | 3100 | -17.2 | 47.6 | 74°10'44.0''S 162°30'53.0''E |
| Mt. New Zealand | 2 | South | 3100 | -17.2 | 47.6 | 74°10'44.0''S 162°30'53.0''E |
| Mt. New Zealand | 3 | South | 3100 | -17.2 | 47.6 | 74°10'44.0''S 162°30'53.0''E |

### Environmental DNA extraction and Denaturing Gel Gradient Electrophoresis

Environmental DNA was extracted from 0.3 g of crushed rocks using NucleoSpin^®^ Plant II Kit (Macherey-Nagel, Gmbh & Co. KG, Duren, Germany) according to the manufacturer’s instructions and quantified by Quant-iT dsDNA HS assay kit (Invitrogen molecular probes-Eugene, Oregon, USA). Microbial diversity was screened by Denaturing Gel Gradient Electrophoresis (DGGE). A semi-nested PCR was performed using primers with a GC-clamp (Muyzer et al., 1993); fungal and algal ITS rRNA was amplified from environmental DNA with primers ITS1F-GC/ITS2 (Gardes and Bruns, 1993; White et al., 1990), following the protocol reported in Selbmann et al. (2017), while 341F-GC/518R primers (Muyzer et al., 1993) were utilized for 16S rRNA amplification (Valášková and Baldrian, 2009) (Table 2).

**Table 2.**
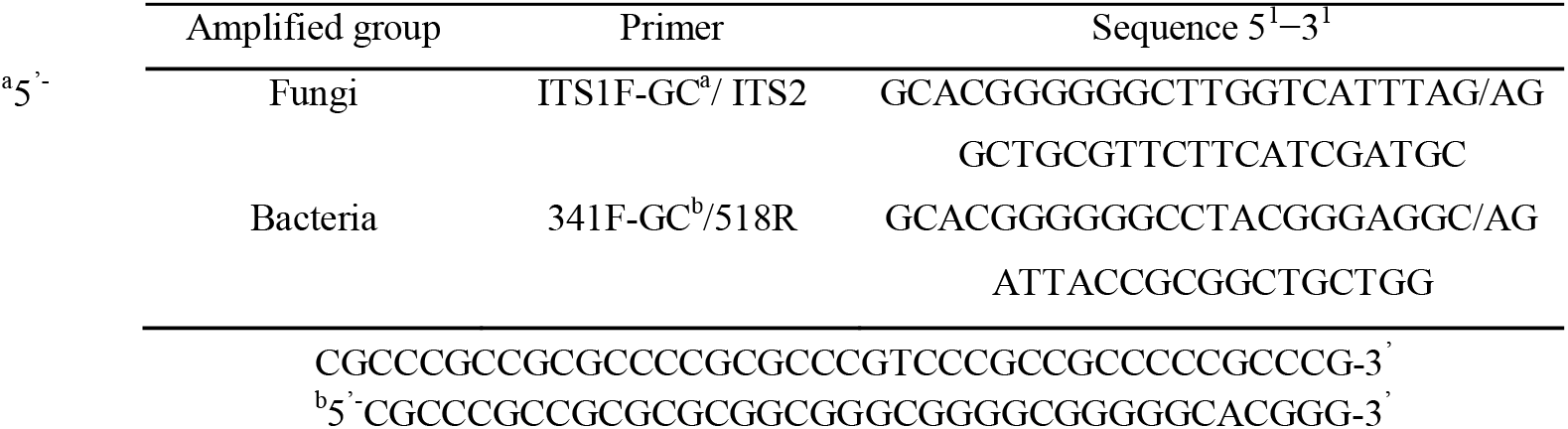
PCR primers used for DGGE in the present study.

Amplicons were purified using NucleoSpin^®^ Gel and PCR Clean-up (Macherey-Nagel, Gmbh & Co. KG, Düren, Germany). One-hundred ng of final DNA concentration were loaded into each well for DGGE and runs were performed on DGGE Electrophoresis System (C.B.S. Scientific, Del Mar, California, USA). Gels at 7.5% polyacrylamide (37.5:1 acrylamide:bisacrylamide) were mixed in a gradient maker (Hoefer, USA), using two different concentrations: for fungi and algae (0% and 70%) and for bacteria (0% and 60%). The electrophoresis run was performed for 5 h for fungi and algae and 3.5 h for bacteria in 1× TAE Buffer at a constant temperature of 60 °C and 200 V.

Bands were visualized by staining for 40 min with GelRed solution (Biotiuminc, CA, USA) (1.34 g NaCl, 66.7 pi GelRed and 200 mL dw) and then visualized with an UV transilluminator (Chemidoc, Bio-Rad). Scanned gels were analyzed with TotalLab Quant Software (Cleaver Scientific Ltd; United Kingdom): bands were assigned and matched automatically and then checked manually. Profile similarity was calculated by determining Dice’s similarity coefficient, considering presence/absence of bands, for the total number of lane patterns from the DGGE gels. The similarity coefficients calculated were then used to generate the dendrograms utilizing the clustering method Unweighted Pair Group Method using Arithmetic mean (UPGMA).

Triplicate samples were processed and analyzed independently.

### NMDS ordination plots

Multivariate statistical analyses were performed to determine the effects of sun exposure on microbial diversity composition using PAST software (PAleontological STatistics, ver. 2.17) The effect of this abiotic parameter was tested displaying changes in communities’ composition with Non-Metric Multidimensional Scaling (NMDS) based both on abundance data (bands intensity), calculating Bray-Curtis distance index and presence–absence data, using Jaccard index (Clarke, 1993). Means of abundance data were square-root transformed and analyses were carried out with 999 permutations as described in Coleine et al. (2018b). NMDS were plotted using the combined occurrence and abundance data of the three replicates from each site. Permutational multivariate analyses of variance (PERMANOVA, p<0.05) based on the Euclidean distance were utilized to establish differences in the two differently sun-exposed rock-inhabiting communities.

### Quantitative PCR

Total fungal and bacterial abundances were measured for all samples by quantitative PCR (qPCR) using NS91F (5’-GTCCCTGCCCTTTGTACACAC-3’) and ITS51R (5’-ACCTTGTTACGACTTTTACTTCCTC-3’) for total fungi and Eub338 (5’-ACTCCTACGGGAGGCAGCAG-3’) and Eub518 (5’-ATTACCGCGGCTGCTGG-3’) for total bacteria (Fierer et al., 2005). To determine relative gene-copies abundances, standard curves were generated using a 10-fold serial dilution of a plasmid containing a copy of *Cryomyces antarcticus* ITS rRNA gene for fungi and *Escherichia coli* 16S rRNA gene for bacteria. The amplicons were generated in 100 μl reactions, containing 50 μl of 2x PCR BioMix™ (Bioline, London, UK), 5 pmol of both forward and reverse primers, 43 μl of DEPC water and 5 μl template of genomic DNA. For fungi, amplification of ITS region was carried out as follows: 95 °C for 3 min and the 35 cycles of 95 °C 40 s, 55 °C 30 s, and final extension at 72 °C 40 s; while bacterial 16S region was amplified as follows: 95 °C for 3 min and the 35 cycles of 95 °C 40 s, 53 °C 30 s, and final extension at 72 °C 40 s. PCR products were then purified using the NucleoSpin^®^ PCR Clean-up kit (MACHEREY-NAGEL, GmbH & Co. KG), quantified with Qubit dsDNA HS Assay kit and cloned using the pGEM^®^-T Easy Vector Systems (Promega, Madison, Wisconsin, US). Plasmids were isolated using the NucleoSpin Plasmid kit (Macherey-Nagel, GmbH & Co. KG).

Five standards were utilized for qPCR in series from 10^7^ to 10^3^ copies.

The 25□μl qPCR reactions contained 12.5□μl iQ™ SYBR^®^ Green Supermix (Bio-Rad, Hercules, California, US), 1□μl of each forward and reverse primers, 0.3 ng of environmental DNA or standard and 9.5□μl nuclease-free water. The reactions were carried out on Quantitative real-time BioRad CFX96™ PCR detection system (Bio-Rad, Hercules, California, US). Primers and amplification protocols are mentioned before. Melting curves were generated to confirm that the amplified products were of the appropriate size. Fungal and bacterial gene copy numbers were generated using a regression equation for each assay relating the cycle threshold (*C*_t_) value to the known number of copies in the standards.

Each assay included no-template controls (NTC). All qPCR reactions were run in triplicate.

Means and standard deviations were calculated and statistical analysis were performed using one-way analysis of variance (Anova) and pairwise multiple comparison procedures, carried out using the statistical software SigmaStat 2.0 (Jandel, USA). Significant differences were calculated by Tukey test (p<0.05).

The fungal-to bacterial ratio was calculated from log-transformed abundance values.

## Results

### DGGE profiles

DNA was efficiently extracted from almost all rock samples, with few exceptions (i.e. Linnaeus Terrace south) where DNA extraction was not quantifiable, but PCR-DGGE worked out for all samples.

Because results were similar when analysing fungi and algae, only data based on fungi are reported.

Profile similarity based only on band presence/absence, was calculated by the Dice coefficient and UPGMA was used to create dendrograms describing pattern similarities (Figs. 1, 2). Overall, the banding patterns of the replicates showed a high degree of similarity (data not shown), which was also supported by the dendrograms, generating DGGE patterns that grouped together as most similar to each other. A coherent grouping according to locations and to sun exposure was generated in the clustering based on fungal, and, even more clearly, on bacterial profiles, with few scattered exceptions. In particular, in the clustering generated on fungal profiles, samples were split according to the localities for University Valley, Pudding Butte, Battleship Promontory, Mt. New Zealand and Finger Mt. (Fig. 1). The grouping was also evident in the clustering based on DGGE bacterial profiles (Fig. 2); as for fungi, the splitting was obtained in the communities from Pudding Butte (south), Battleship Promontory (south) and Finger Mt. (south), but also for Linnaeus Terrace and Knobhead for both sun expositions.

**Figure 1.**
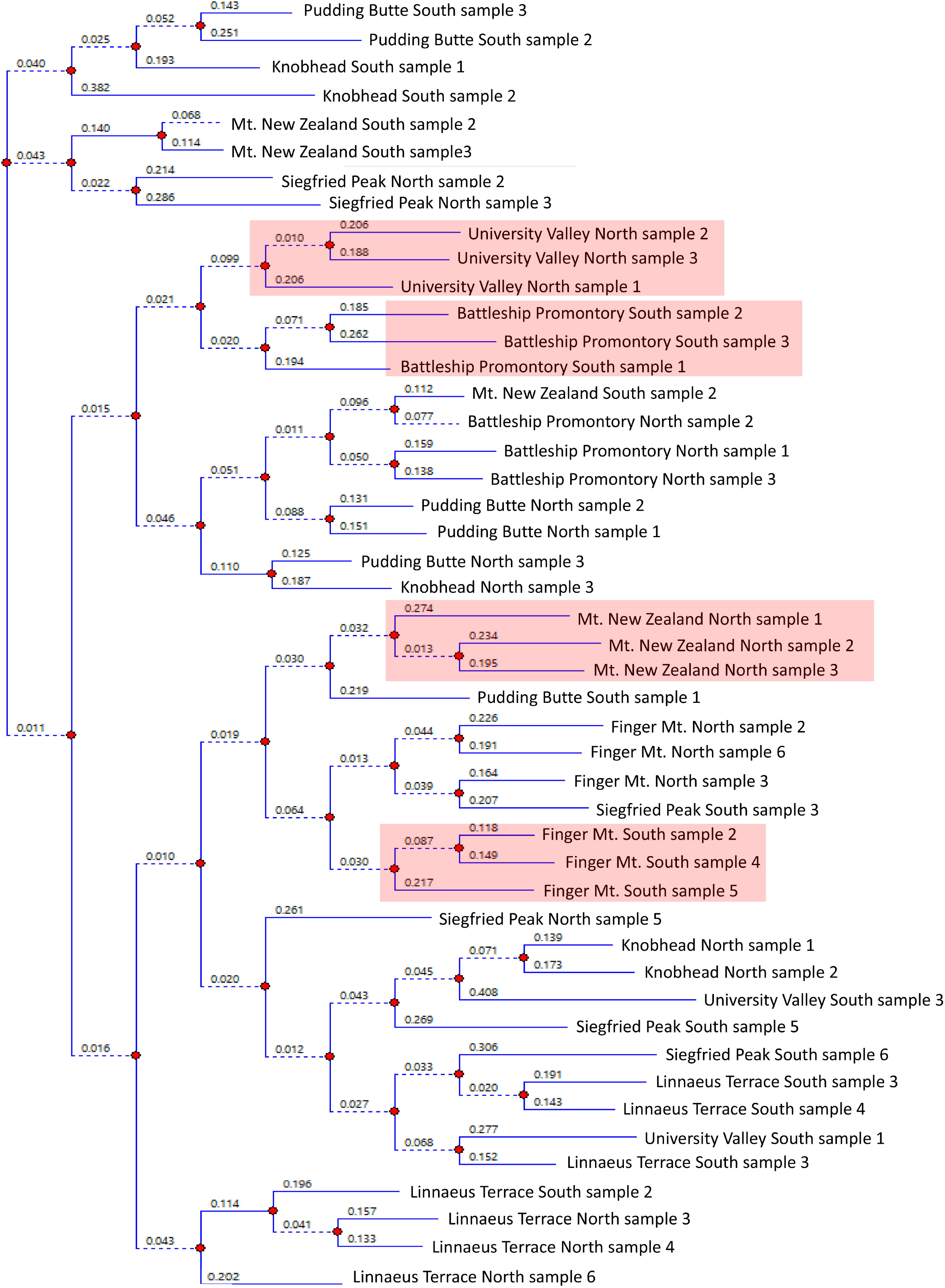
Dendrogram obtained by Unweighted Pair Group Method using Arithmetic average (UPGMA) based on the DGGE profiles of the fungal component of the endolithic communities. The relationships among samples are based on similarity, evaluated by the Dice coefficient. Triplicate samples were analysed.

**Figure 2.**
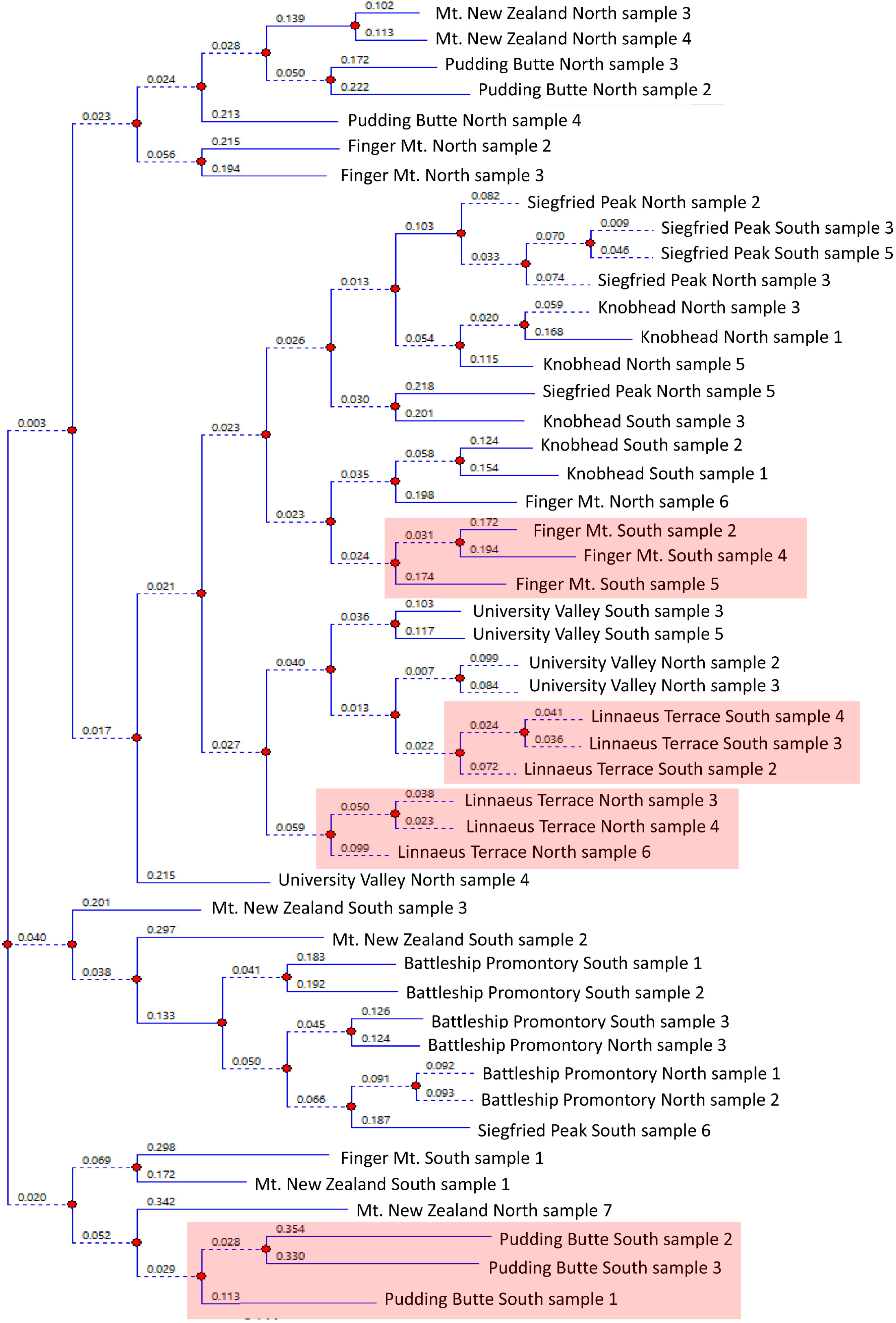
Dendrogram obtained by Unweighted Pair Group Method using Arithmetic average (UPGMA) based on the DGGE profiles of the bacterial component of the endolithic communities. The relationships among samples are based on similarity, evaluated by the Dice coefficient. Triplicate samples were analysed.

### NMDS ordination plots

To investigate the similarity of the fungal and bacterial communities’ composition amongst different sun exposures, a Non-metric Multi-Dimensional Scaling (NMDS) analysis was computed.

NMDS ordination plots were generated both with the only presence-absence matrix using the Jaccard index and with the combined frequency of occurrence using the Bray-Curtis index. Because both approaches produced similar results, we showed results based on abundance only.

When stress values are <0.1, the NMDS plot is considered to be an acceptable representation of the original data; in fact, a stress value below 0.1 indicates a reliable ordination of data, without a real probability of misinterpretation (Clarke, 1993). In this analysis, the stress value was 0.09 for fungi and 0.07 for bacteria, fitting with the ideal ordination.

NMDS plots generated from the DGGE profiles of amplified fungal ITS and bacterial 16S rDNA revealed that only small changes occurred among samples collected in the same sun-exposed rock surface (p>0.05) and did not exhibit any changes endolithic communities by sampled localities (data not shown). On the contrary, a major change (1-way NPMANOVA, p<0.05) occurred between north and south sun-exposed communities, showing a strong structuring of fungal and bacterial communities according to the sun exposure (Fig. 3).

**Figure 3.**
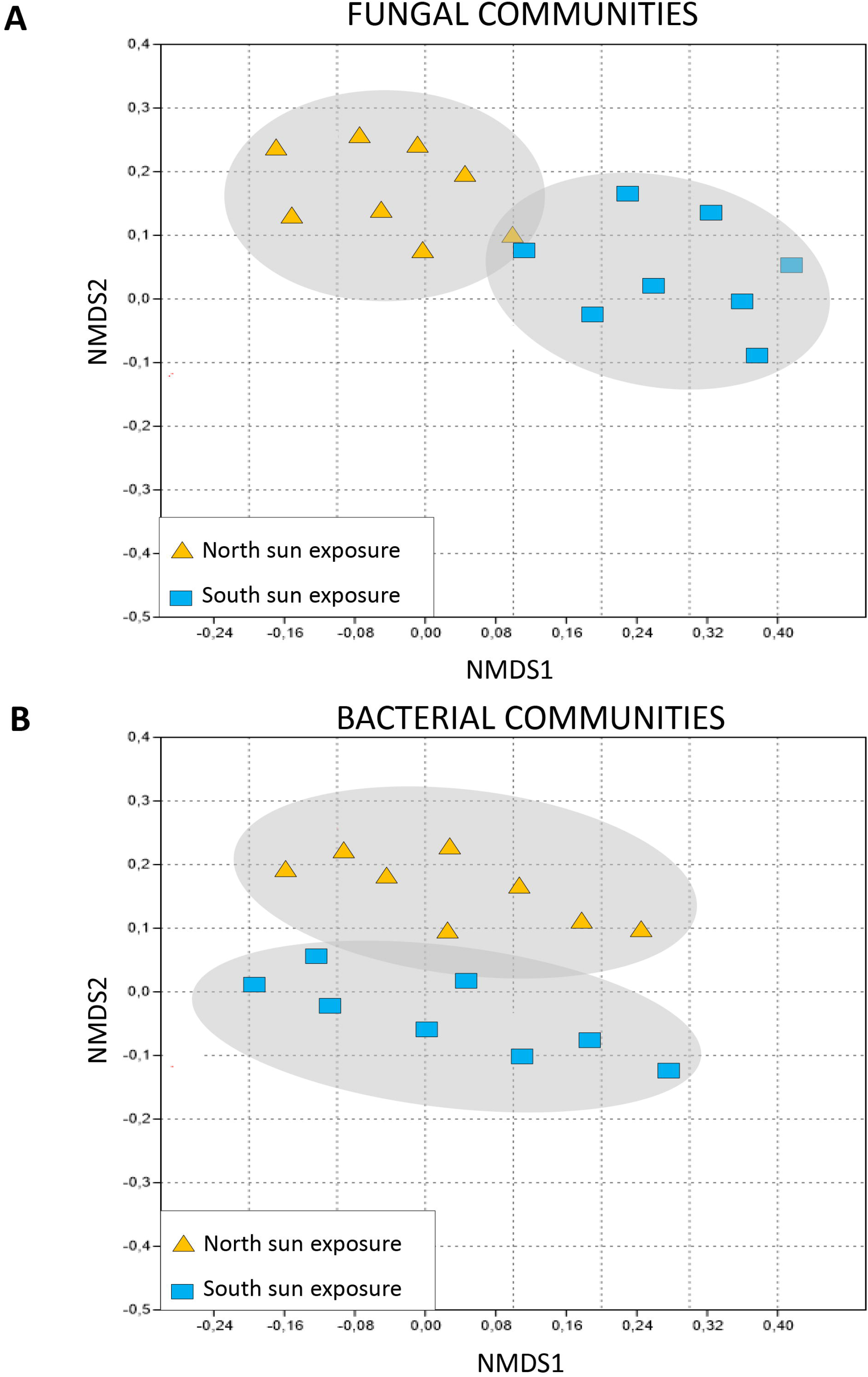
Non-Metric Multidimensional Scaling (N-MDS) ordination plots for fungal (A) and bacterial endolithic communities (B), calculating the Bray–Curtis index, based on square-root transformed abundance data. Stress value (A): 0.09. Stress value (B): 0.07

### Abundance of fungal and bacterial communities

Both ITS and 16S rRNA gene copy numbers varied mostly significantly (p <0.05) along the sampled sites, ranging from 6.1 × 10^4^ (Knobhead south sun exposure, Dry Valleys, Southern Victoria Land) to 9.8 × 10^6^ fungal copies (Battleship Promontory south sun exposure, Dry Valleys); conversely, the differently north and south exposed rock surfaces of Linnaeus Terrace (Dry Valleys) showed the highest (7.7 × 10^5^) and the lowest (1.3 × 10^3^) bacterial gene-copies, respectively (Fig. 4 and Table 1S).

**Figure 4.**
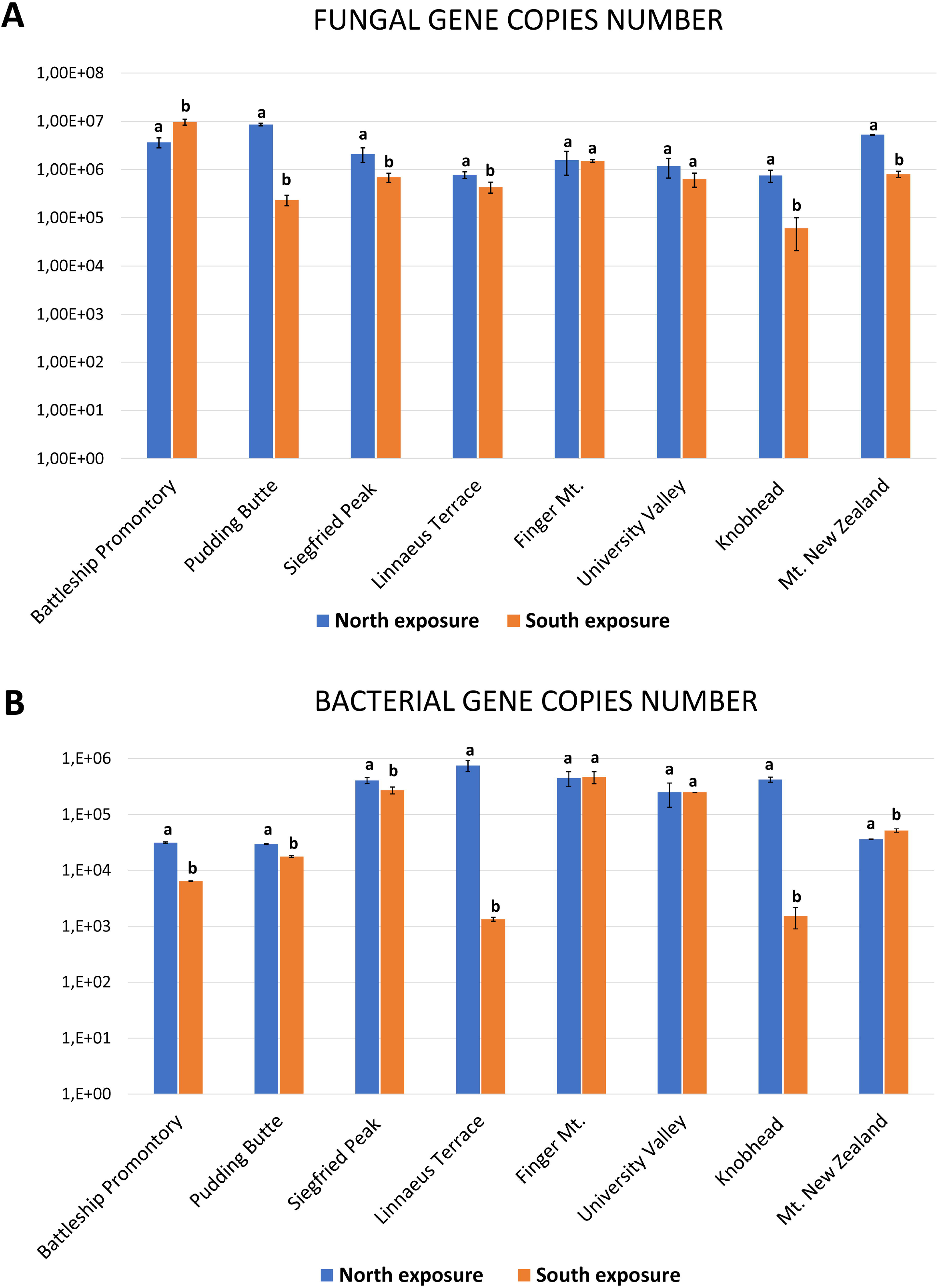
Abundances of total fungi and bacteria in endolithic communities, as estimated using the qPCR assays. Error bars are the standard errors. Significant differences are calculated by Tukey test with p value<0.05 and indicated by different letters.

Overall, both fungal and bacterial abundances varied between the two sun-exposures, except in the case of Finger Mt. and University Valley in Dry Valleys, where abundance of the two major biological compartments was similar. Furthermore, microbial abundance was generally higher in north-exposed rocks, with only few exceptions: fungi were most abundant in south sun-exposed samples at Battleship Promontory, while bacteria were larger in number at Mt. New Zealand south (Fig.4).

The fungal-to bacterial ratio (F/B) (based on log-copy numbers) showed a slightly higher dominance of fungi in these ecosystems in both sun-exposed sampled localities and were significantly different in all samples (p<0.05). In north-exposed rocks F/B was 1.24±0.2 (mean ± SD), while in south-exposed surface it varied between 1.37 ± 0.31 (data not shown).

## Discussions

The Antarctic cryptoendolithic communities host among the most resistant microorganisms on Earth, being able to survive under environmental conditions once accounted as incompatible with life (Horowitz et al., 1972).

Recent studies have improved knowledge of the microbial diversity and composition of these ecosystems (Wei et al., 2016; Archer et al., 2017) and provided important insights to clarify the complex relationship between environmental parameters (i.e. altitude and distance from sea) and biodiversity (Selbmann et al., 2017; Coleine et al., 2018a, b). The lack of consistent microbial diversity patterns along altitudinal and sea distance gradients has suggested that new hypotheses about which can act as driving factor shaping biodiversity must put forward (Selbmann et al., 2017; Coleine et al, 2018a).

Towards this end, the aim of this study was to test the potential effects on the structure and composition of the bacterial, fungal and algal assemblages of a new parameter, namely sun exposure, as significant variable influencing endolithic settlement and development, using a preliminary molecular screening approach on a considerable amount of rock samples collected along the Victoria Land (Continental Antarctica).

We found that DGGE-based similarity assessment visualized as dendrograms indicated a clear grouping of samples according mostly by sun exposure, highlighting an evident relationship between microbial composition and this environmental parameter. This trend is consistent in all biological compartments examined. Particularly, a clear clustering was observed for DGGE profiles obtained from samples collected in Pudding Butte, Battleship Promontory, Mt. New Zealand and Finger Mt. in fungi and algae (Fig. 2, data not shown); as for eukaryotes, in bacteria, clear groups were observed in samples belonging to Battleship Promontory, Finger Mt., Pudding Butte, Knobhead and Linnaeus Terrace.

Considering all the data, higher similarities in fingerprinting profiles were observed within samples collected in the same site, with very few scattered exceptions, likely due to variability among samples.

These findings highlighted the effectiveness of sampling strategy applied for the Antarctic Expedition 2015-16 and did not reflect what was observed by Selbmann et al. (2017). In that study, based on a previous sampling (PNRA, Antarctic Expedition 2010-11), the largest Antarctic sampling to date of rocks hosting lithic communities, including different rock typologies (i.e. sandstone, granite, dolerite) from 46 different localities in Victoria Land, was investigated. Results indicated a remarkable local variability found even in rocks from the same site, maybe due to the size of sampled area (about 100 mq^2^) and to variability of rock typologies collected.

DGGE band profiles were analysed by statistical analysis and, consistently to clustering analysis, samples were mainly grouped by sun exposure along the three analysed biological groups. The observed shift in community composition correlated to sun exposure was also confirmed by Non-Metric Multidimensional scaling (NMDS) analysis, which organizes data into 2-D spatial graphs by reducing dimensionality. An apparent gradient in abundance and community composition in response to sun exposure was previously observed in Coleine et al. (2018b), where authors investigated biodiversity and composition of functional groups of fungi in Antarctic endolithic communities and reported the absence of any correlation with altitude and sea distance, while a remarkable variability was observed considering the sun exposure parameter.

It was previously hypothesized that the distribution of endoliths reflects the degree of insulation on the rock surfaces; in northern exposed rocks, environmental conditions are more favourable than southern exposed faces, and cryptoendolithic colonization is more often observed and favoured (Friedmann, 1977; Friedmann and Weed, 1987). The capability to maintain biological activity may depend on sufficient insulation of the rock to allow an efficient photosynthetic process; moreover, warmer temperatures will allow metabolic activity and more water due to snow melt (McKay and Friedmann, 1985; Deegenaars and Watson, 1998).

Our findings support the hypothesis that sun radiation not only affects temperature of rocks surface, availability of water and metabolic activity, but even the biodiversity.

Community structure was also shaped by north and south exposure when fungal and bacterial abundances were estimated with a qPCR assay. In the most localities, fungal (ranging from 6.1 × 10^4^ to 9.8 × 10^6^ gene copies) and bacterial (7.7 × 10^5^ to 1.3 × 10^3^ gene copies) abundances changed significantly according to sun exposure.

We also calculated the fungal:bacterial (F/B) ratio, a metric to assess environmental impacts and the functional implications of microbial communities (Raeymaekers, 2000; Fierer et al., 2005). The F/B dominance was significantly affected by sun exposure and showed a significative dominance of fungi in these microbiomes. Fungi predominate in the south-exposed rocks (with the only exception of Finger Mt. in McMurdo Dry Valleys) where conditions are much more extreme; indeed, we found a greater fungal dominance (F/B, 1.37) respect than north-exposed sites (F/B, 1.24).

We propose that further studies are needed to elucidate the relationship between fungi and bacteria, and to reveal their functions in these ecosystems. Nevertheless, this result is not surprisingly: extreme environments (i.e. the ice-free desert of Victoria Land in Continental Antarctica) are not an only prerogative of archaea and bacteria. Among eukaryotes, fungi (alone or in symbiosis with cyanobacteria or algae forming lichens) are the most versatile and ecologically successful phylogenetic lineage; they evolved to survive and proliferate in the extremes (Magan, 2007), even in habitats normally precluded to the most (Selbmann et al., 2013). Fungi are, for instance, generally more resistant to desiccation than bacteria with hyphae that may cross air-filled soil pores to access nutrients and water (Gordon et al., 2008; de Vries et al., 2012), showing a remarkable ability to survive in stress conditions (e.g. low water availability) of Antarctic ice-free areas (Onofri et al., 2004).

The most remarkable example for stress resistance is, in fact, given by the fungus *Cryomyces antarcticus*, a cryptoendolithic black fungus isolated from the McMurdo Dry Valleys in Antarctica, chosen as the best eukaryotic test organisms for astrobiological investigations for its stunning resistance to temperature cycles (−20/+20 °C), high temperature (+90 °C) and saline concentration (up to 25% NaCl) (Onofri et al., 2008; 2012; 2015). It was found to resist to ionizing radiation up to 55.81 kGy (Selbmann et al., 2018), while the bacterium *Deinococcus radiodurans*, widely considered the extremophile par excellence and gold-medalist of radiation resistance (Battista et al., 1997; Venkateswaran et al., 2000), survives up to 20 kGy of gamma radiation.

In conclusion, even though DGGE-based approach does not show individual taxa responses and gives limited insight into taxonomy, it is still currently considered a powerful, rapid and costly-effective method for determining shapes in microbial community composition (Zheng et al., 2013; Kovalski Mitter et al., 2018). This technique gave us the advantage to rapidly and inexpensively analyze a great number of rock samples and three biological domains (fungi, algae and bacteria), providing a clear trend of how sun exposure influences cryptoendolithic community biodiversity and composition.

This study expands existing knowledge on the relationship between environmental parameters and Antarctic endolithic biodiversity and examines endolithic community structure. The results inform planning of future Antarctic campaigns to maximize identification of active endolithic communities.

With the advent of -omics approaches such as metabarcoding, metagenomics and metabolomics, we may explore the composition deeply focusing on a targeted selection of rock samples. These samplings will characterize taxa identity and abundance in rocks hosting endolithic communities, examine the complex community dynamics, their stress-adaptation strategies, and potential functions across an environmental variation gradient that is influenced by sun exposure.

A detailed picture on how these communities respond to increasing environmental pressures will also give clues for predicting and monitoring the effects of global change on these unique border ecosystems and their vulnerable biodiversity.

## ACKNOWLEDGMENTS

L.S., C.C. and L.Z. wish to thank the Italian National Program for Antarctic Researches (PNRA) for funding sampling campaigns and researches activities in Italy in the frame of Projects 2009/A1.11, 2013/AZ-17, 2015/AZ1.02 and AMunDsEN PNRA_00006. The Italian Antarctic National Museum (MNA) is acknowledged for financial support to the Mycological Section on the MNA for preserving rock Antarctic samples analysed in this study and stored in the Culture Collection of Fungi from Extreme Environments (CCFEE), University of Tuscia, Italy.

---

**Tables 1S.**
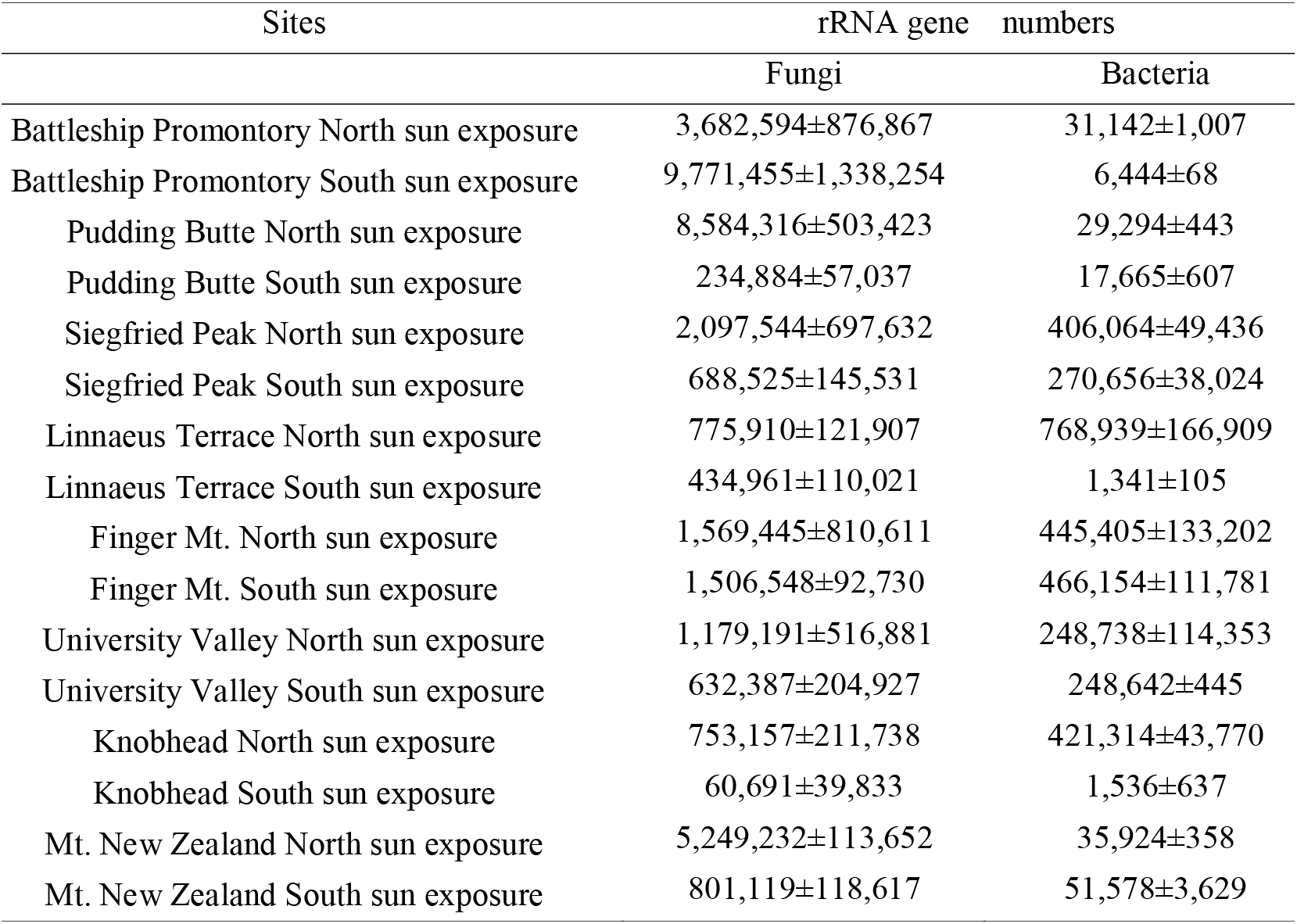
Fungal and Bacterial gene-copies abundances. Mean and deviation standard are listed in for each site.

